# Vertical Stratification in Urban Green Space Aerobiomes

**DOI:** 10.1101/2020.06.28.176743

**Authors:** Jake M. Robinson, Christian Cando-Dumancela, Craig Liddicoat, Philip Weinstein, Ross Cameron, Martin F. Breed

## Abstract

Exposure to a diverse environmental microbiome is thought to play an important role in ‘educating’ the immune system and facilitating competitive exclusion of pathogens to maintain human health. Vegetation and soil are known to be key sources of airborne microbiota––the *aerobiome*. Only a limited number of studies have attempted to characterise the dynamics of the aerobiome, and no studies to date have investigated these dynamics from a vertical perspective simulating human exposure. Studies of pollution and allergenic pollen show vertical stratification at various scales, and present an expectation that such vertical stratification may also be present in the aerobiome. Such stratification could have important implications for public health and for the design, engineering and management of urban green spaces. For example, do children receive the same exposure to airborne microbiota as taller adults, and what are the downstream implications for health? In this study, we combine an innovative columnar sampling method at soil level, 0.0, 0.5, 1.0, and 2.0 m together with high-throughput sequencing of the bacterial 16S rRNA gene to assess whether significant vertical stratification of the aerobiome occurred in a parkland habitat in Adelaide, South Australia. Our results provide evidence of vertical stratification in both alpha and beta (compositional) diversity of airborne bacterial communities, with diversity increasing roughly with height. We also found significant vertical stratification in known pathogenic and beneficial bacterial taxa, suggesting potentially different exposure attributes between adults and children. These results could have important implications for public health and urban planning, potentially informing ways to optimise the design and management of health-promoting urban green spaces.

## 1 Introduction

Over the last 100 years, urban populations have increased dramatically (Cox et al. 2018). Indeed, urbanisation is predicted to increase further with an estimated 60% of the world’s population living in towns and cities by 2030 (Hake et al. 2016; Dogan and Gurcan, 2019). The ecological footprint associated with increased urbanisation has exerted considerable pressure on both local and planetary systems (While and Whitehead, 2013; Maheshwari et al. 2020). Moreover, a growing body of evidence now links urbanisation to the rise in noncommunicable diseases (NCDs) such as chronic inflammatory conditions (e.g., autoimmune disorders and allergies) and the transmission of communicable diseases such as dengue fever, chikungunya and the recent COVID-19 outbreak (Andrea, 2019; Goryakin et al. 2017; Alirol et al. 2011; Ali and Dasti, 2018; Franchi, 2020; Wu et al. 2020).

Biodiversity loss is a global megatrend, with current species extinction rates estimated to be 1,000 times higher than historical background rates, and future rates likely to increase to 10,000 times higher (Haahtela et al. 2013; De Vos et al. 2015). This is driven in part by urbanisation, and associated processes including unsustainable land use, resource exploitation, pollution and climate change (Sol et al. 2014; Hughes, 2017; Crenna et al. 2019). Importantly, the biodiversity loss and NCD global megatrends are thought to be interrelated (Von Hertzen et al. 2015; Haahtela, 2019).

Intensive land-use and reduced biodiversity at the macro-scale (e.g., substrates, plant communities and animals) is associated with reductions in biodiversity and structural changes at the micro-scale (i.e., the microbiome) (Bender et al., 2016; Heiman et al. 2016; Blum et al. 2019; Liddicoat et al. 2019); yet microbiomes can be restored through revegetation (Gellie et al. 2017; Mills et al. 2020). These changes could have implications for human health as exposure to a diverse suite of environmental microbes is important to ‘educate’, regulate and maintain the human immune system (Rook et al. 2003; Rook et al. 2013; Arleevskaya et al. 2019). Furthermore, studies now link the microbiome to a plethora of maladies from Alzheimer’s disease and myalgic encephalomyelitis, through inflammatory bowel and skin diseases, to respiratory health (Hansom and Giloteaux, 2017; Prescott et al. 2017; Sokolowska et al. 2018; Aschard et al. 2019; Kowalski and Mulak, 2019).

Environmental factors are thought to be more important than genetic factors in shaping the composition of the gut microbiome (Rothschild et al. 2018). Indeed, prior research suggests that early life exposure to a diverse range of environmental microbes is particularly important (until the weaning age––typically 0-4 years); during this period the composition of the human gut microbiome is highly dynamic and readily colonised by environmental microbes (Yang et al. 2016; Moore and Townsend, 2019). However, recent research suggests the adult microbiome exhibits greater plasticity than previously thought. For example, Martinson et al. (2019) provided evidence to suggest that certain bacterial families in the adult human gut microbiome, such as *Enterobacteriaceae*, exhibit high levels of colonisation plasticity. Furthermore Schmidt et al. (2019) recently showed that one in three microbial cells from the oral environment pass through the digestive tract to settle and replenish the gut microbiome of healthy adult humans. Browne et al. (2016) showed that spore-forming bacteria (which survive in aerobic conditions) dominate the human gut, comprising 50-60% of bacterial genera, and display greater change in abundance and species over time compared to non-spore formers, suggesting that many gut bacteria may come and go from the environment. This presents a challenge to both the notion of an oral-gut barrier (Martinsen et al. 2005) and the level of microbiome stability in adulthood (D’Argenio and Salvatore, 2015; Stearns et al. 2017). Is it therefore possible that exposure to environmental microbes remains important throughout the life-course (e.g., for immunoregulation, competitive exclusion of pathogens, and homeostasis)?

Vegetation and soil are known to be key sources of airborne microbiota––i.e., the *aerobiome* (Joung et al. 2017; Liu et al. 2018). A small number of studies have attempted to characterise the community structure and spatiotemporal dynamics of the aerobiome. For example, Mhuireach et al. (2016) compared bioaerosol samples in green spaces and parking lots and found compositional distinctions in bacterial communities between the two land cover types. Furthermore, Mhuireach et al. (2019) explored spatiotemporal controls on the aerobiome and suggested that localised site factors were likely to be important in driving bacterial community structure. However, no known studies have investigated the spatial and compositional factors from a vertical perspective, simulating potential human exposures at different heights.

Support for the existence of aerobiome stratification can be drawn from studies of pollution, allergenic pollen and fluid dynamics of particulates where vertical stratification has been shown to occur at various scales. For example, in an internal environment and under ventilated conditions, Miles (2008) showed that NH3 molecule concentrations decreased vertically with increasing distance from source (i.e., the ground). Gou and Nui (2007) found that vertical concentration stratification of particles up to PM10 (10.0μm) occurred under different ventilation conditions. Particles smaller than 2.5μm were less affected by gravitational factors, and submicron particles with small relaxation times (i.e., the time required for particles to adjust their velocity to new conditions of forces) behaved more like trace gases following main airstreams. Alcázar et al. (1998) found higher concentration of *Urtica membranacea* pollen at the upper region of their sampling height range of 1.5 m-15 m, and higher concentrations of *U. urens-Parietaria sp*. at lower heights––possibly due to pollen mass and different fluid dynamics.

The existence of aerobiome vertical stratification could have important implications for the design, engineering and management of urban green spaces––particularly those aimed at promoting public health via microbial exposure (Watkins et al. 2020). For example, do children receive the same exposure to airborne microbiota as taller adults? Do people who lie down or work close to the ground (e.g., gardeners bending over to dig) have different exposure levels to those who remain upright, and what are the downstream implications for health? Developing a refined understanding of this aerobiome-human interface could also have implications for the design and monitoring of nature-based health interventions, for example via green/nature prescribing (Robinson and Breed, 2019; Shanahan et al. 2019; Robinson et al. 2020).

Protocols for sampling the aerobiome to date have often included a reasonable yet arbitrary sampling height of 2 m (Airaudi et al. 1996; Cordeiro, 2010; Mhuireach et al. 2016; Domingue, 2017). Therefore, investigating aerobiome composition at various heights could provide important methodological insights to fine-tune future study protocols and public health recommendations. In this proof of concept study, we combine innovative columnar aerobiome sampling methods along with remote sensing techniques and high-throughput sequencing of the bacterial 16S rRNA gene. The primary objectives of this study were to: (a) assess whether significant vertical stratification in bacterial species richness and evenness (alpha diversity) of the aerobiome occurred; (b) assess whether significant compositional differences (beta diversity) between sampling heights occurred; and (c) to preliminarily assess whether there were significant altitudinal differences in known pathogenic and beneficial bacterial taxa.

## 2 Materials and Methods

### 2.1. Site selection

Our study site comprised three vegetated plots totalling seven ha of the southern section of the Adelaide Parklands (Kaurna Warra Pintyanthi), South Australia. The justification for the selected study site was as follows:

1. Its broadly consistent soil geochemistry, as the southern Parklands generally fall within the Upper Outwash Plain soil boundary (coalescing alluvial soil, draining the Eden Fault Block).
2. This area is managed by a single division of the City of Adelaide, minimising variation in site management and allowing for simpler study logistics.
3. A single study site (i.e., the southern section) in the Parklands provided a degree of control over potential variation in landscape effects on the aerobiome (e.g., dominant vegetation type, distance to coast, elevation, orientation, aspect).
4. Urban Parkland is representative of conditions that both child and adult residents might be exposed to.

Following site selection, boundaries of three plots (as polygons) were defined in QGIS 3 (v3.0.2). These polygons were subsequently converted to shapefiles (.shp) and a random point algorithm was generated. This provided randomly selected sampling points within each vegetated plot to include in our study (Figure 1). The spatial coordinates for each sampling point were recorded and programmed into a handheld global positioning system (GPS) device. This was operated on site to allow us to identify the relevant locations for setting up the sampling stations.

**Figure 1.**
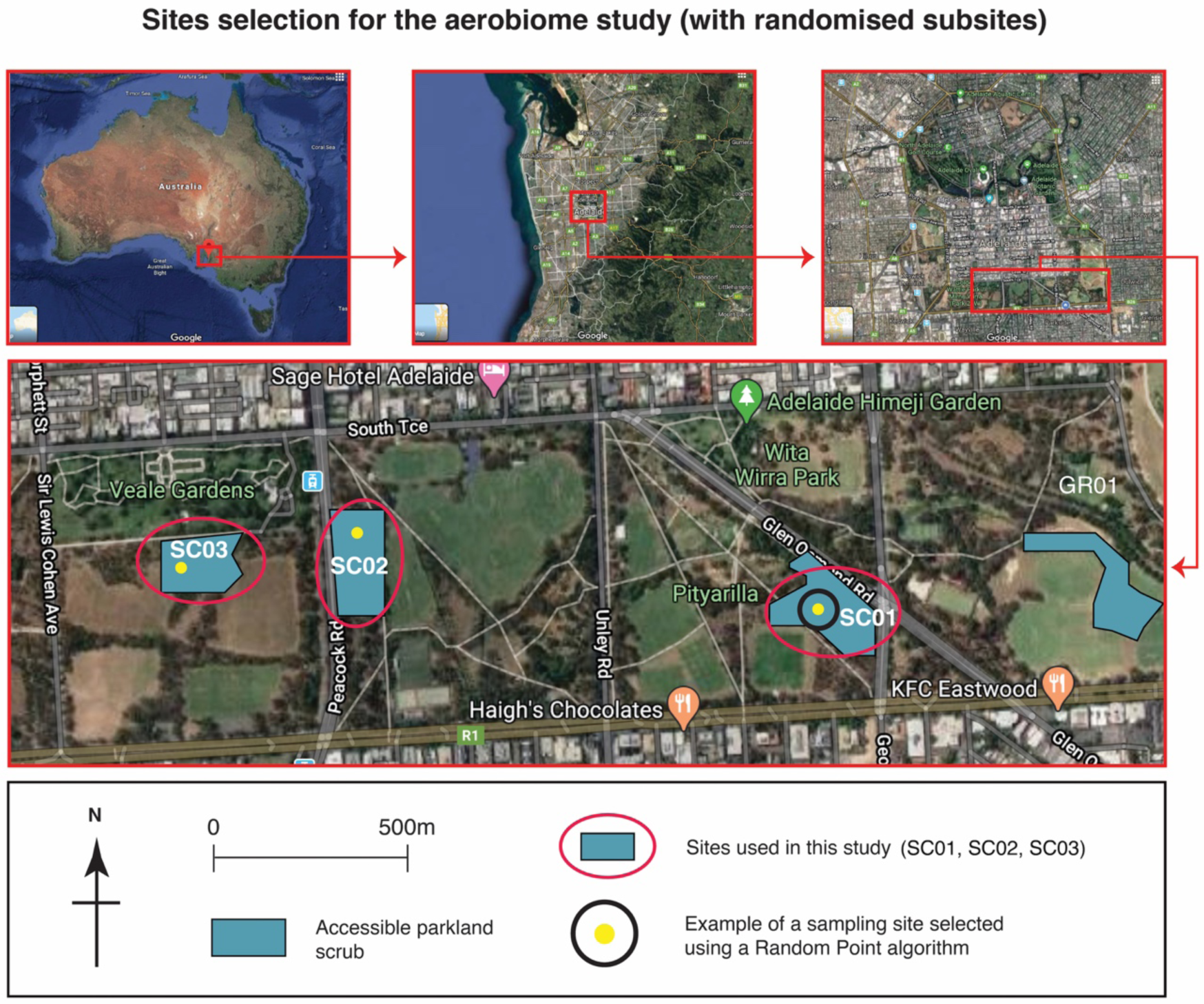
Location of study sites, showing the randomly selected sampling locations.

### 2.2. Sampling equipment

The sampling stations (Figure 2) were constructed using timber (SpecRite 42 mm x 28 mm x 2.7 m screening Merbau). The sampling stations comprised a timber stand with 45° leg braces. Hooks and guy ropes were also installed, ensuring stability in the field. Steel brackets were installed to secure petri dishes, which we used to passively sample the aerobiome as per Mhuireach et al. (2016).

**Figure 2.**
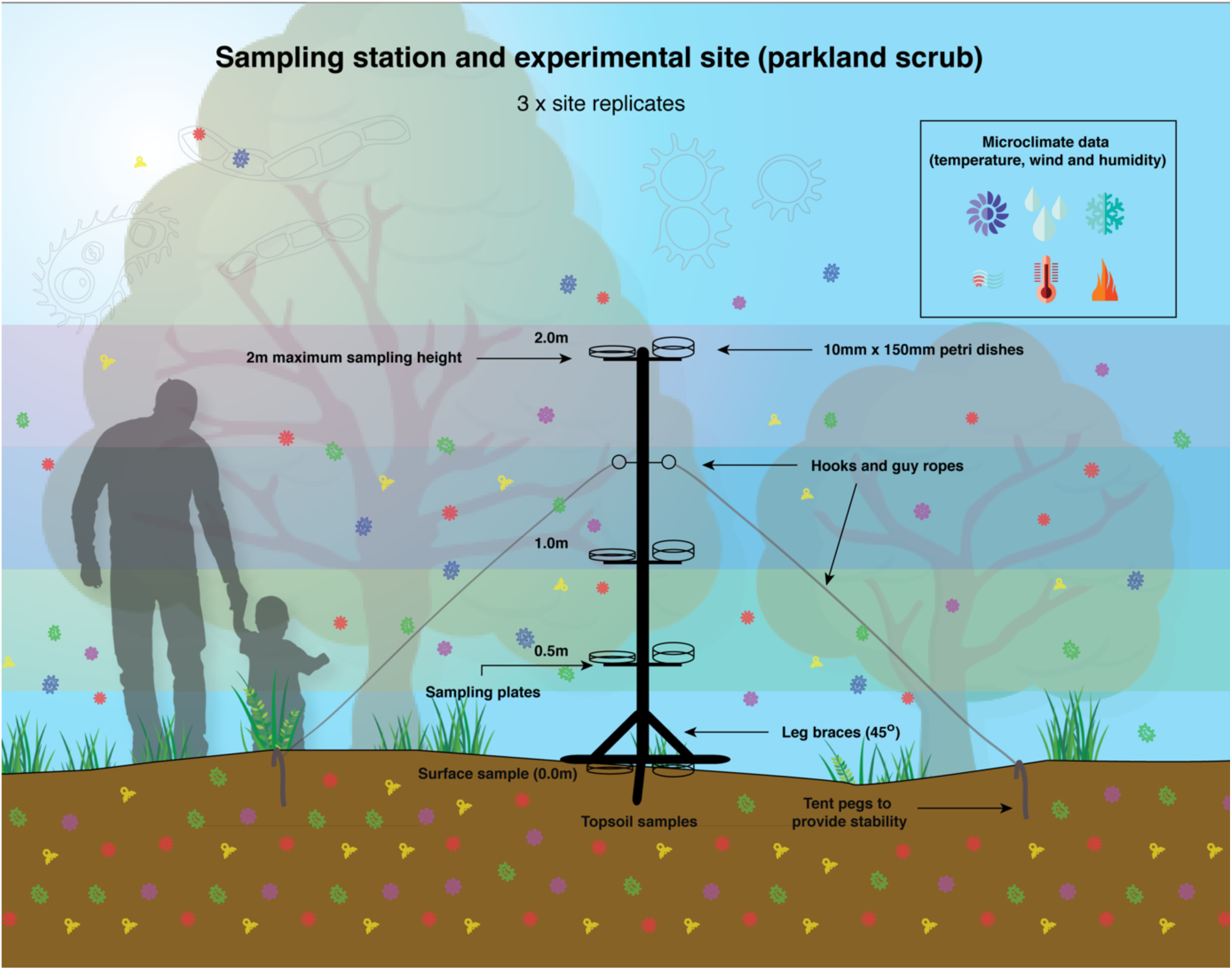
Design of the aerobiome vertical stratification sampling stations. These were installed in scrub habitat in the Adelaide Parklands.

The level of stability was tested in two phases – *Phase 1*: during windy conditions (∼Beaufort scale No. 5) in a yard environment, and *Phase 2*: in situ, prior to the sampling phase.

### 2.3. Data loggers

We installed temperature and relative humidity data loggers at each sampling station. Each logger was programmed to record data at 8-second intervals for the entire sampling period. The dataloggers were calibrated using a mercury thermometer and a sling psychrometer.

### 2.4. On-site setup procedure

The sampling stations were placed into position between 0600-0800hrs on 4th, 5th and 6th November 2019. This ensured sufficient time was allocated to travel between the sampling locations. From 0800hrs onwards and prior to installing the petri dishes for passive sampling, the sampling stations were decontaminated using a 5% Decon 90 solution. The microclimate data loggers were then decontaminated and installed on the sampling stations. The nearest trees (all <10 m height and 20 cm-50 cm in diameter at breast height) were between 2 m and 5 m from the sampling stations.

### 2.5. Sampling protocol

The sampling procedure involved collecting soil samples (actively) and airborne microbiota (passively). Environmental metadata were also collected (e.g., windspeed, temperature and relative humidity). Soil pH at each site was measured using a digital pH meter (Alotpower). The probe of the pH meter was inserted into the soil and left for a period of 1-minute prior to taking a reading, as per manufacturer’s instructions.

Windspeed and direction data for the entire study area were obtained from Adelaide’s meteorological weather station at Ngayirdapira (West Terrace): Lat: −34.93, Lon: 138.58, Height: 29.32 m. Windspeed and direction was also recorded at each sampling site on an hourly basis (Mhuireach et al. 2016) using the handheld anemometer (Digitech *QM-1644)*.

#### 2.5.1. Soil samples

Topsoil samples were collected using a small shovel and stored in 50 mL sterile falcon tubes. The shovel was decontaminated using the 5% Decon 90 solution prior to use. Wearing gloves, we sampled five topsoil samples (depth: 5-7cm) at equidistant sampling points, 20-30 cm from the central stem of each sampling station (Zarraonaindia et al. 2015). The soil samples were subsequently pooled and then homogenised, passed through a 1 mm pore sieve, and placed in new sterile 50 mL Falcon tubes. The sample tubes were labelled using a predefined labelling system. We included field controls of soil samples by opening 50 mL sterile falcon tubes for 60 s at each site (Mbareche et al. 2019). All soil and field control samples were immediately chilled by placing in an ice box in the field, and then storing at −80°C in the lab prior to DNA extraction and sequencing (Zarraonaindia et al. 2015). In total, we collected 15 soil subsamples per sampling day across the three sampling stations for each of the three sampling days. Subsamples were pooled and homogenised by sampling station and day, which gave a total of nine homogenised samples (three per sampling station) plus three field controls.

#### 2.5.2. Aerobiome samples

Passive sampling methods were used to collect low biomass aerobiome samples following established protocols (Mhuireach et al. 2016; Mhuireach et al. 2019). Petri dishes (100 × 15 mm) were attached with decontaminated Velcro tabs on the sampling stations at four sampling heights: ground level, 0.5 m, 1 m, and 2 m. The total height of the sampling stations was 2 m from ground level (95% of adult human heights lie within 2 SD at 1.93 m). One metre is the average height of a 4-year old child (typically the maximum weaning age––and the time when the gut microbiome is thought to become less plastic), the height of a pram bassinet, and the height of an adult sitting in a chair (Dettwyler 2017; Milani et al. 2017; RCPCH, 2020). Fifty cm is the approximate height of a pushchair seat, and of an adult torso and head (representing the height of an adult sitting on the floor). The surface is also an important sampling level, for example, representing the point of contact for a crawling child or an adult lying on the floor. The steel petri dish sampling plates were also decontaminated using the 5% Decon 90 solution prior to use.

The petri dishes were secured to the sampling stations (Figure 2) and were left open for 6-8 hours (Mhuireach et al. 2016). At the end of the sampling period, we closed the petri dishes. A new set of gloves was worn for the handling of petri dishes at each vertical sampling point to reduce contamination. The petri dishes were then sealed using Parafilm, labelled, immediately placed on ice, and transported to the laboratory for storage at −80°C prior to DNA extraction (Mhuireach et al. 2019). Unused petri dishes were left open for 60 s, and then sealed at each site as field controls. Dishes were later swabbed during the DNA extraction process using nylon flocked swabs (FLOQSwabs Cat. No. 501CS01, Copan Diagnostics Inc., CA, USA) (Mhuireach et al. 2019; Bae et al. 2019; Liddicoat et al. 2020).

### 2.6. DNA extraction, amplification and sequencing

We extracted DNA from samples at the Evolutionary Biology Unit (EBU), South Australian Museum. The order of processing samples was randomised using a digital number randomiser, including the soil samples (higher biomass), which were processed after the low biomass, aerobiome samples to minimise cross-contamination.

The petri dishes for each sampling station were swabbed with FLOQSwabs for 30 s (with consistent back and forth strokes) in a laminar flow cabinet type 1 (License No. 926207). The base and lid samples for each height, station and date were then pooled, prior to extraction. The swabs were cut with decontaminated scissors directly into labelled 2 mL Eppendorf tubes. We used Qiagen QIAamp DNA Blood Mini Kits to extract DNA from the swabs together with extraction blank controls, and Qiagen DNAeasy PowerLyzer Soil Kits to extract DNA from the soil samples (and extraction blank controls). We followed the manufacturer’s instructions throughout the extraction process.

PCR amplification was done in triplicate using the 341F/806R primer targeting the V3-V4 region of the 16S rRNA gene (5’ -CCTAYGGGRBGCASCAG-3’/5’ - GGACTACNNGGGTATCTAAT-3’). The 300 bp paired end run was sequenced on an Illumina MiSeq platform at the Australian Genome Research Facility Ltd (AGRF) using two flowcells (ID 000000000-CW9V6 and 000000000-CVPGT). Image analysis was done in real time by the MiSeq Control Software (MCS) v2.6.2.1 and Real Time Analysis (RTA) v1.18.54. Then the Illumina bcl2fastq 2.20.0.422 pipeline was used to generate the sequence data.

### 2.7. Bioinformatics and statistical analysis

Paired-end reads were assembled by aligning the forward and reverse reads using PEAR (version 0.9.5). Primers were identified and trimmed. Trimmed reads were processed using Quantitative Insights into Microbial Ecology (QIIME 1.8.4), USEARCH (version 8.0.1623), and UPARSE software. Using USEARCH tools, reads were quality filtered, full length duplicate reads were removed and sorted by abundance. Singletons or unique reads in the data set were discarded. Reads were clustered and chimeric reads were filtered using the “rdp_gold” database as a reference. To obtain the number of reads in each operational taxonomic unit (OTU), reads were mapped back to OTUs with a minimum identity of 97%. Taxonomy was assigned using QIIME.

We used the phyloseq package (McMurdie and Holmes, 2013) in R to import and analyse the sequencing data, and decontam (Davis et al. 2018) to identify and exclude contaminants. Lower biomass samples (i.e., air, field blanks, and extraction blank controls) were analysed using the isNotContaminant() function, where contaminants were identified by increased prevalence in negative controls. Higher biomass samples (i.e., soil, and corresponding extraction blanks) were analysed using the isContaminant() function. Using isContaminant(), contaminants were identified by the frequency that varies inversely with sample DNA concentration, or by increased prevalence in negative controls. All taxa identified as contaminants were pooled and removed from further analysis. To estimate OTU alpha diversity we derived Shannon Index values based on rarefied abundances (Liddicoat et al. 2019) in phyloseq. We generated box and violin plots with ggplot2 (Wickham and Wickham, 2007) to visualise the distribution of the alpha diversity scores for each sampling height. Microbial beta diversity was visualised using non-metric multidimensional scaling (NMDS) ordination of Bray-Curtis distances based on rarefied OTU abundances. The ordinations plots show low-dimensional ordination space in which similar samples are plotted close together, and dissimilar samples are plotted far apart.

We used permutational multivariate analysis of variance (PERMANOVA) to test for compositional differences between sampling heights. The Pearson’s product-moment and Spearman’s rank correlation tests were used to examine correlations between sampling height and alpha diversity scores. A Mann-Whitney Wilcoxon test was used to examine differences in alpha diversity between merged air sampling heights (0.0 −0.5 m and 1.0-2.0 m) and a Kruskal Wallace chi-squared test to explore differences in correlations between sites and dates. We also calculated OTU relative abundances using the phyloseq package in R to examine the distribution of taxa that have potential implications for public health. To compare presence and proportions of taxa we used 2-sample tests for equality of proportions with continuity corrections and created radial charts using pivot tables with comma separated value (csv) files.

## 3 Results

We observed a significant negative correlation between alpha diversity (air and soil for all sites/dates) and sampling height (*r* = −0.58, df = 38, *P* = <0.01; Figure 3A; Table 1). Alpha diversity ranged from 1 to 6 and was highest at soil level followed by the lower air sampling levels (0.0 m-0.5 m) and the upper sampling levels (1.0 m-2.0 m), respectively.

**Table 1.**
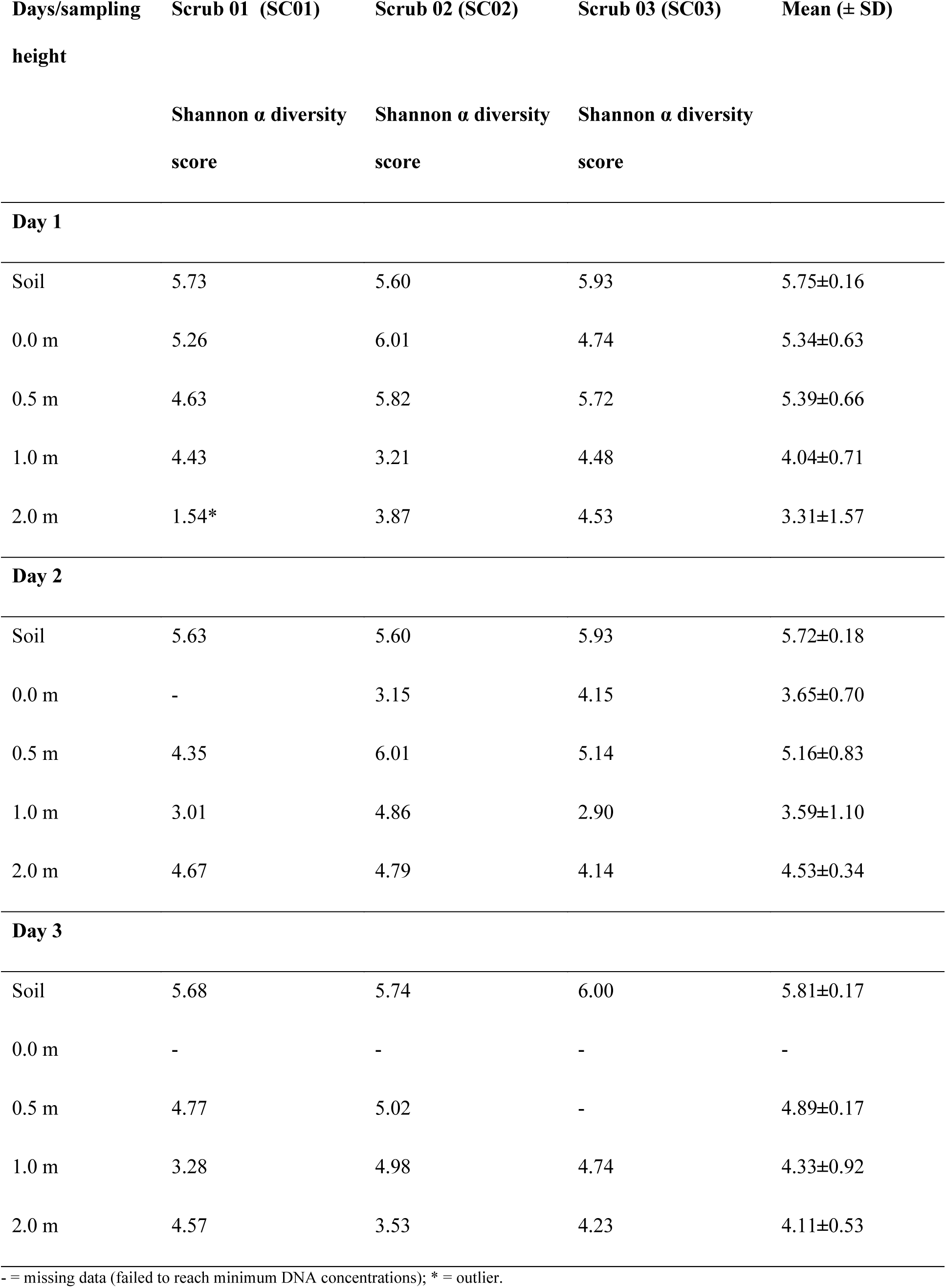
Shannon alpha diversity scores for each spatial and temporal replicate, along with means and standard deviations.

**Figure 3.**
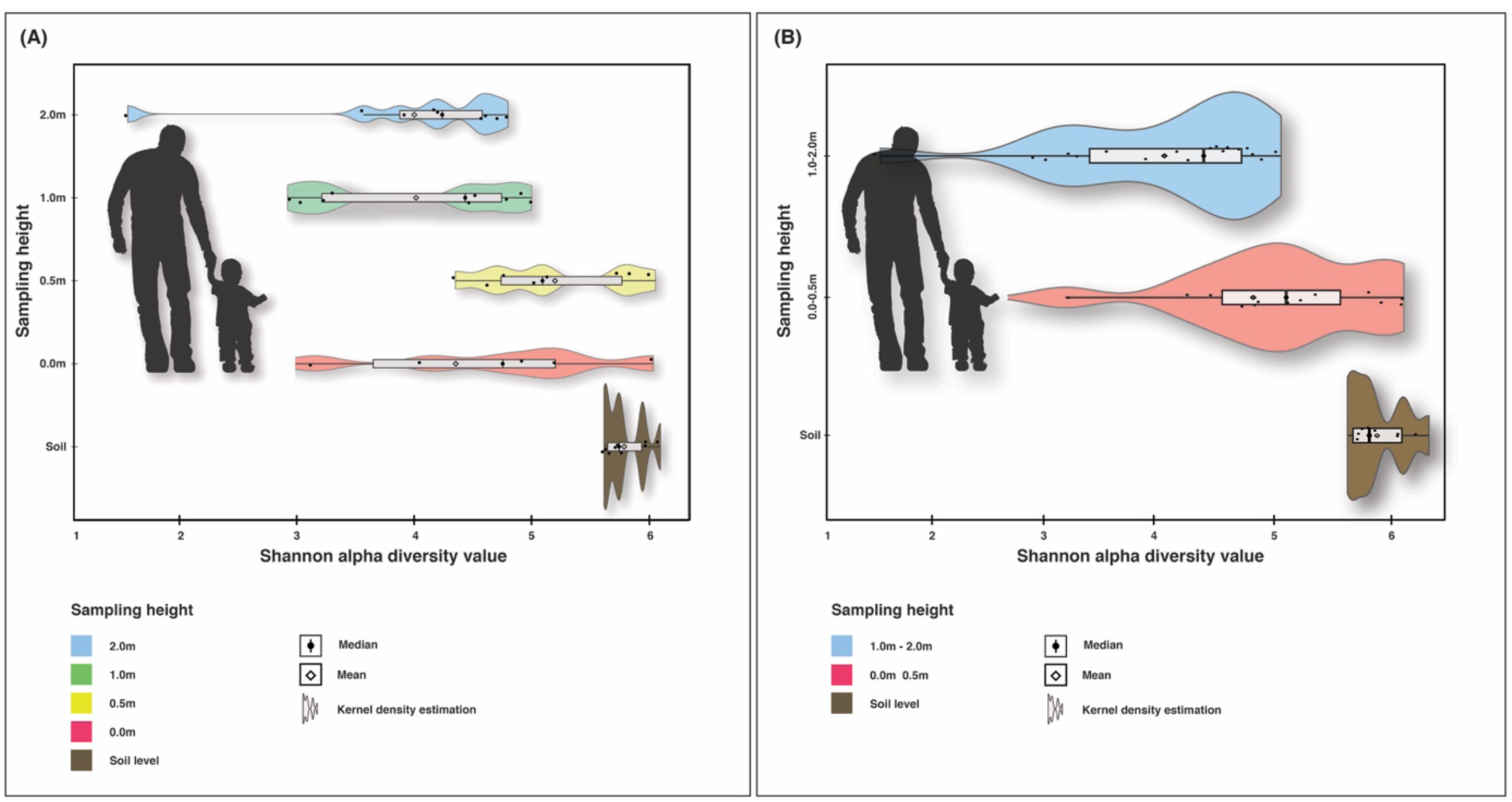
Box/violin plots of Shannon alpha diversity scores for each sampling height including soil (A) and for merged lower heights 0.0-0.5 m and upper heights 1.0-2.0 m, with soil (B). Plots also display mean values, interquartile range and kernel density estimation.

When the lower sampling heights and the upper sampling heights were merged (0.0 with 0.5 m; 1.0 m with 2.0 m), we observed a significant negative correlation between alpha diversity and sampling height (*r* = −0.68, df = 38, *P* = <0.01) (Figure 3B). Following an examination of alpha diversity scores for individual sites and dates, all variants showed negative correlations between alpha diversity and sampling height. Four out of six indicated strong and significant relationships (Day 1: *r* = −0.76, *P* = 0.00; Day 3: *r* = −0.64, *P* = 0.01; SC01: *r* = −0.68, *P* = <0.01; and, SC03: *r* = −0.73, *P* = 0.01; Table 2).

**Table 2.** Pearson’s correlation scores of alpha diversity and sampling height based on all air and soil samples, followed by merged air sampling heights (0.0m-0.5m and 1.0m-2.0m) and soil samples. Correlation scores for each sampling date and site are included.

| Days/sites | *r* score | *df* | *P*-value |
| --- | --- | --- | --- |
| Day 1 (04-11-19) | -0.76 | 11 | <0.01*** |
| Day 2 (05-11-19) | -0.31 | 12 | 0.17 |
| Day 3 (06-11-19) | -0.64 | 11 | 0.01** |
| Scrub 01 (SC01) | -0.68 | 13 | <0.01*** |
| Scrub 02 (SC02) | -0.41 | 12 | 0.14 |
| Scrub 03 (SC03) | -0.73 | 9 | 0.01** |
| *Merged air sampling heights (0.0m-0.5m and 1.0m-2.0m):* |  |  |  |
| Day 1 (04-11-19) | -0.76 | 11 | <0.01*** |
| Day 2 (05-11-19) | -0.59 | 12 | 0.02* |
| Day 3 (06-11-19) | -0.72 | 11 | <0.01*** |
| Scrub 01 (SC01) | -0.72 | 13 | <0.01*** |
| Scrub 02 (SC02) | -0.54 | 12 | 0.04* |
| Scrub 03 (SC03) | -0.86 | 9 | <0.01*** |

With the merged sampling heights, all correlations increased in strength and were all statistically significant (Table 2). A Mann-Whitney Wilcoxson test for differences in alpha diversity between the merged air sampling heights (0.0m-0.5m and 1.0m-2.0m) showed a statistically significant difference (*W* = 188, *P* = <0.01). A Kruskal Wallace chi-squared test indicated no significant difference in correlations between sites or dates (*P* = 0.44).

Using these same merged sampling heights, a 2-sample test for equality of proportions with continuity correction showed a significant difference in proportions of taxa that occurred in lower air sampling heights (compared to upper sampling heights) that also occurred in the soil samples. The positive relationship between the proportion of taxa occurring in the air that also occurred in the soil decreased as vertical distance from the soil increased. For example, at the genus level, 84.4% of taxa in the lower air samples also occurred in the soil samples, whereas only 76.1% of the taxa in the upper air samples occurred in the soil. This difference was statistically significant (Chi-squared = 9.5376, df = 1, *P* = <0.01; Figure 4 shows taxonomic breakdown).

**Figure 4.**
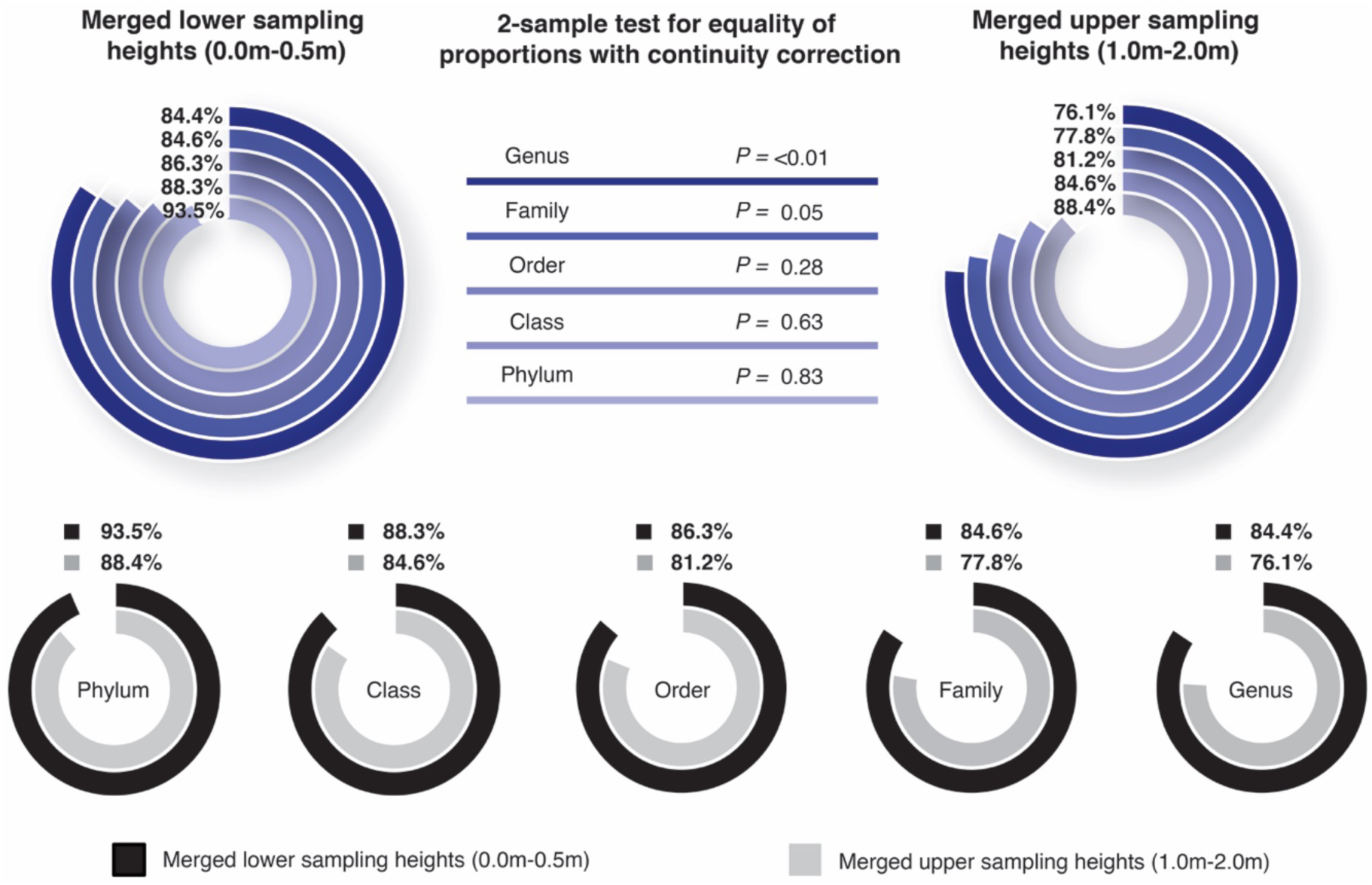
Radial charts showing proportions (as %) of taxa from the air samples that also occurred in the soil samples for each sampling height and across all available taxonomic levels. A 2-sample test for equality of proportions shows significant differences between lower and upper sampling heights for both genus and family taxonomic levels.

Sampling heights displayed distinct bacterial signatures (Figure 5, panel A). Sampling height explained 22% of the variation in environmental microbiota when all air sampling heights and the soil level were included, and this was statistically significant (PERMANOVA df = 4, F = 2.50, R^2^ = 0.22, *P* = <0.01, permutations = 999).

**Figure 5.**
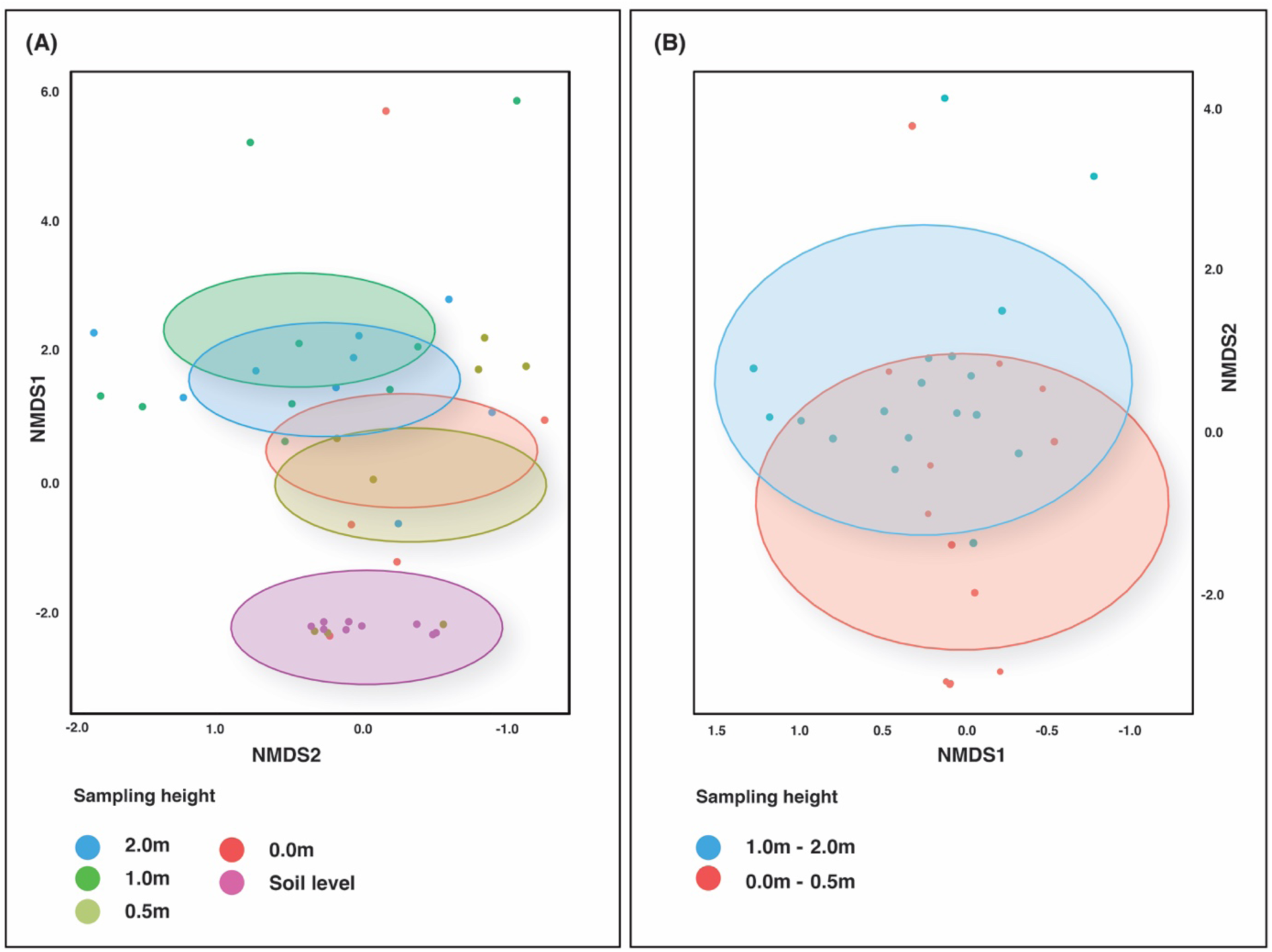
NMDS ordination of bacterial communities for all sampling heights, including soil (A) (Stress: 0.09) and for all sampling heights, excluding soil and merging within lower and upper samples (B) (Stress: 0.10). Ellipses represent Euclidian distance from the centre.

When analysing air samples in isolation, sampling height explained 11% of the variation in environmental microbiota, however, this was not significant (df = 3, F = 1.18, R^2^ = 0.11, *P* = 0.15, permutations = 999). When we merged within lower and upper sampling heights, sampling heights explained 6% of the variation and this was statistically significant (df = 1, F = 1.98, R^2^ = 0.06, *P* = 0.01, permutations = 999) (Figure 5, panel B).

The dominant taxa in the soil and lower sampling heights were Actinobacteria (based on mean relative abundance >1%), and the dominant taxa in the upper sampling heights were Proteobacteria (Figure 6; segments 1 and 9). A significantly greater proportion of Actinobacteria were present in lower air sampling heights (merged 0.0m-0.5m; 43.52% and 26.61%, respectively; 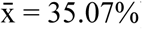) compared to upper air sampling heights (merged 1.0m-2.0m; 17.52% and 19.67%, respectively; 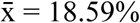) (Chi-squared = 6.1032, df = 1, *P* =0.01).

**Figure 6.**
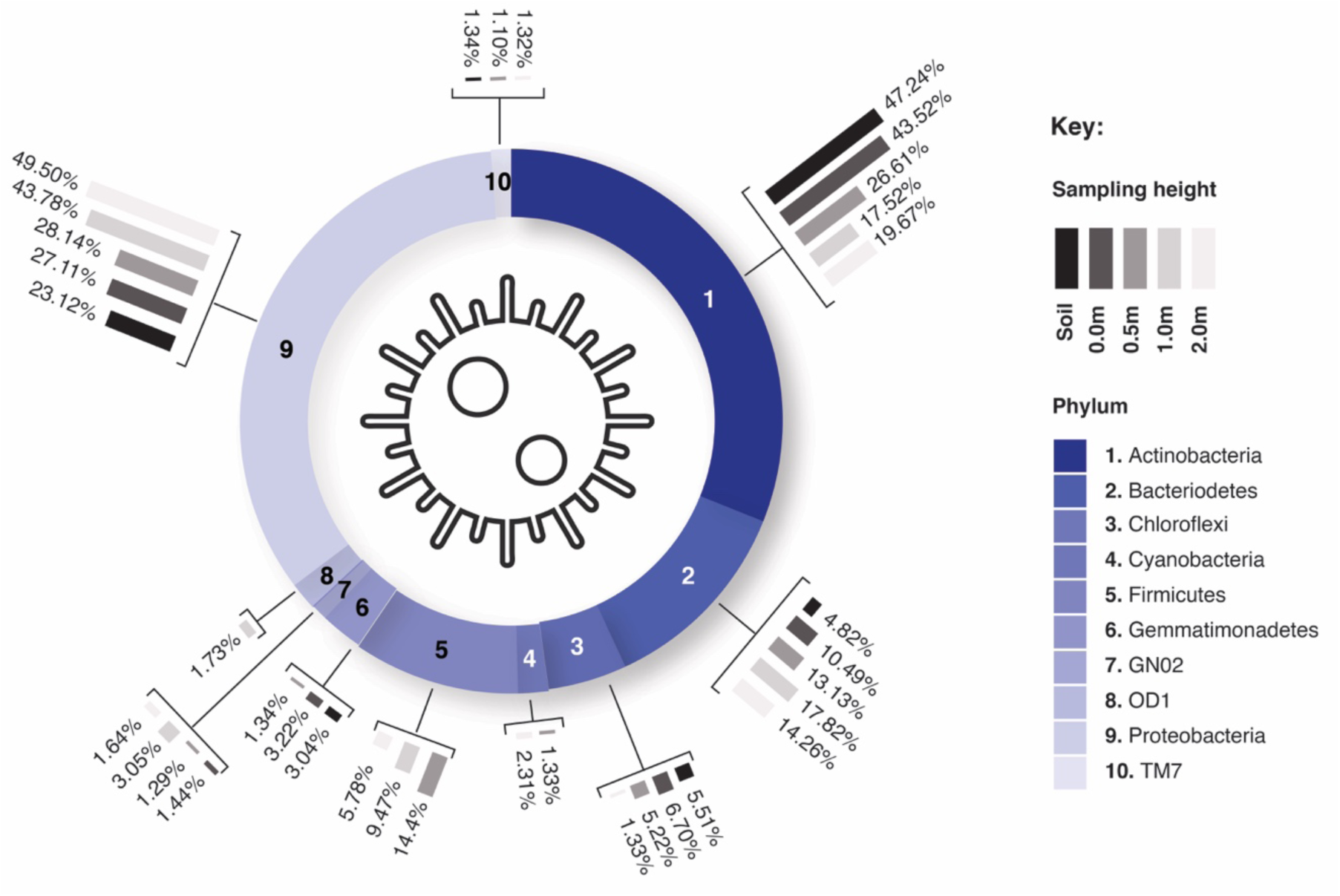
Relative abundance of bacterial OTUs at the phylum taxonomic level (based on mean relative abundance >1% for each sampling height). Ring segments relate to phyla; segment size corresponds to mean relative abundance across all heights; mini bar charts relate to relative abundance of taxa for individual sampling heights where applicable. Actinobacteria (1) dominate lower sampling heights, Proteobacteria (9) dominate upper sampling heights.

**Figure 7.**
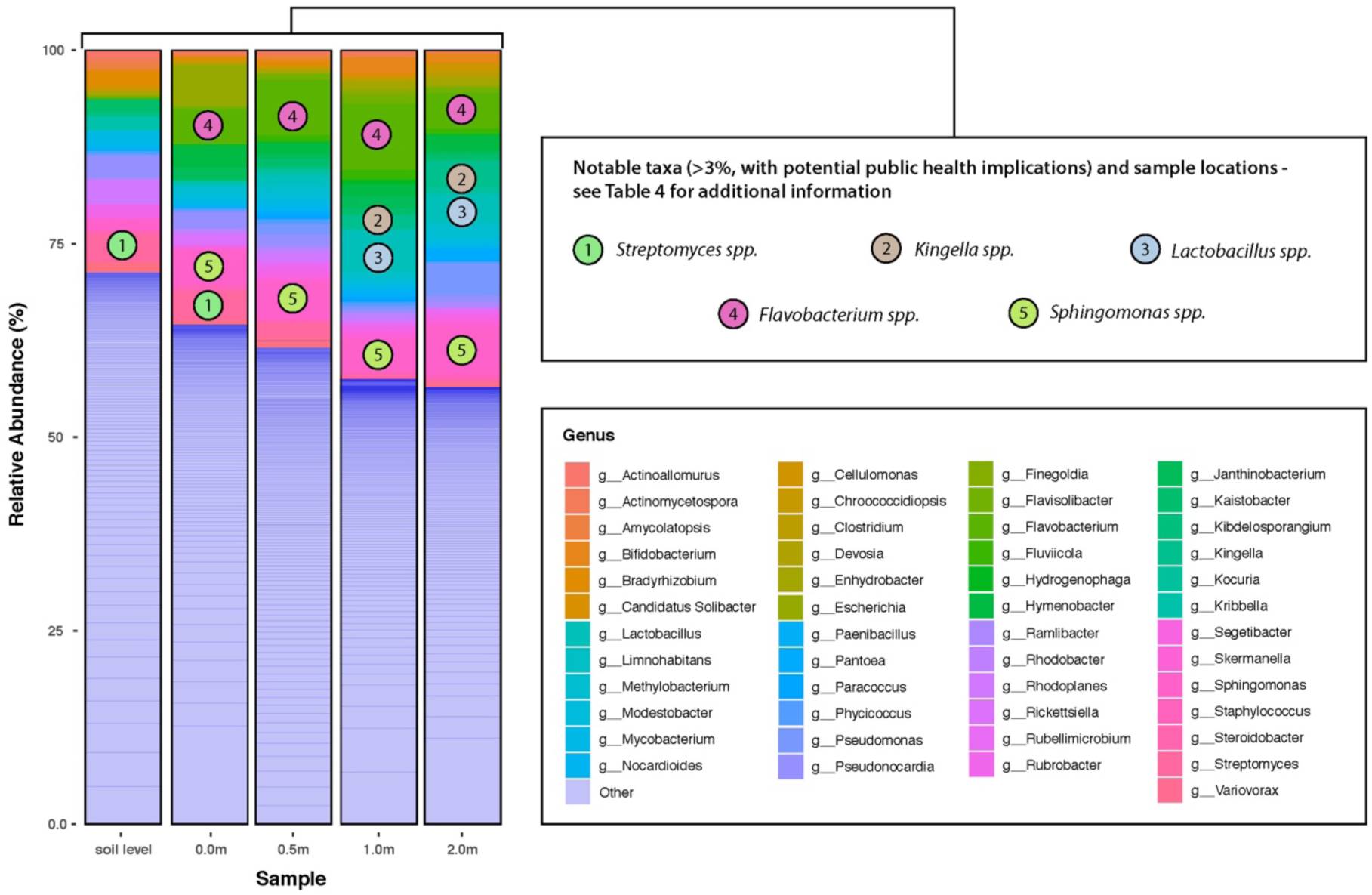
Relative abundance of bacterial OTUs at the genus taxonomic level and identification of notable taxa. Refer to Table 4 for potential public health implications of notable taxa.

A significantly greater proportion of Proteobacteria was present in the upper air sampling heights (merged 1.0m-2.0m; 43.78% and 49.50% respectively; 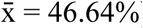) compared to the lower air sampling heights (merged 0.0m-0.5m; 27.11% and 28.14%, respectively; 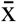 = 27.63%) (Chi-squared = 6.9471, df = 1, *P* = <0.01).

A number of relatively abundant and notable taxa (contingent primarily on their implications for public health) were identified in the samples. The relative abundance of these taxa differed across sampling heights and all significantly correlated with sampling height, ranging from moderate to strong relationships (Table 3) These taxa and their potential implications for public health are highlighted further in Table 4 in the Discussion.

**Table 3.** Spearman’s correlations for notable taxa at the genus level across sampling heights, based on mean relative abundance (>1%) for each sampling height.

| Number | Taxa (genus) | $r_s$ score | S | P-value |
| --- | --- | --- | --- | --- |
| 1 | *Streptomyces* | -0.66 | 23596 | <0.01*** |
| 2 | *Kingella* | +0.39 | 8606 | <0.01*** |
| 3 | *Lactobacillus* | +0.54 | 6470 | <0.01*** |
| 4 | *Flavobacterium* | +0.53 | 6639 | <0.01*** |
| 5 | *Sphingomonas* | +0.39 | 8577 | <0.01*** |

**Table 4.**
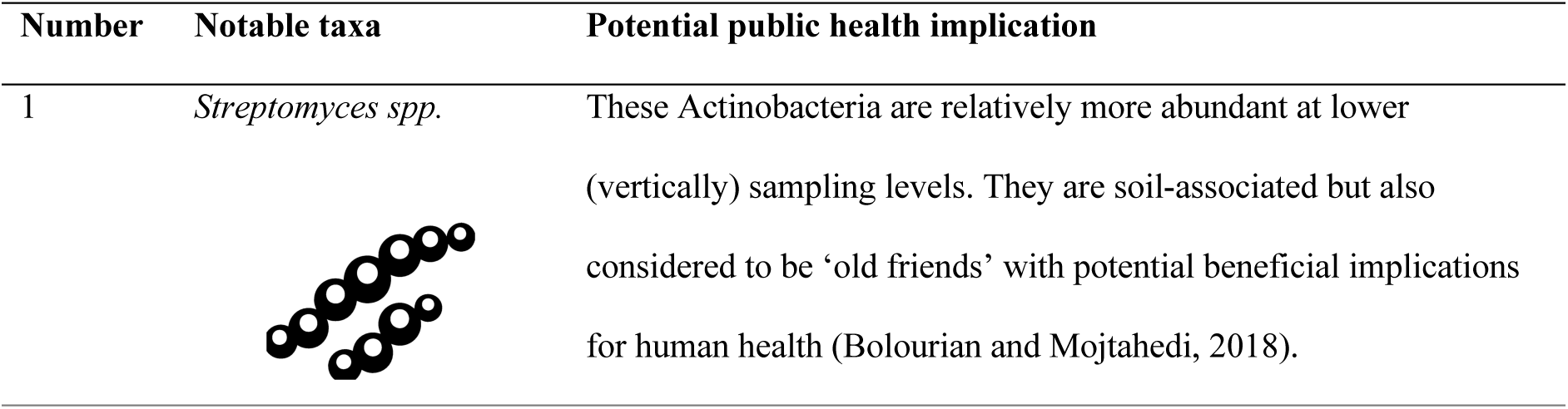
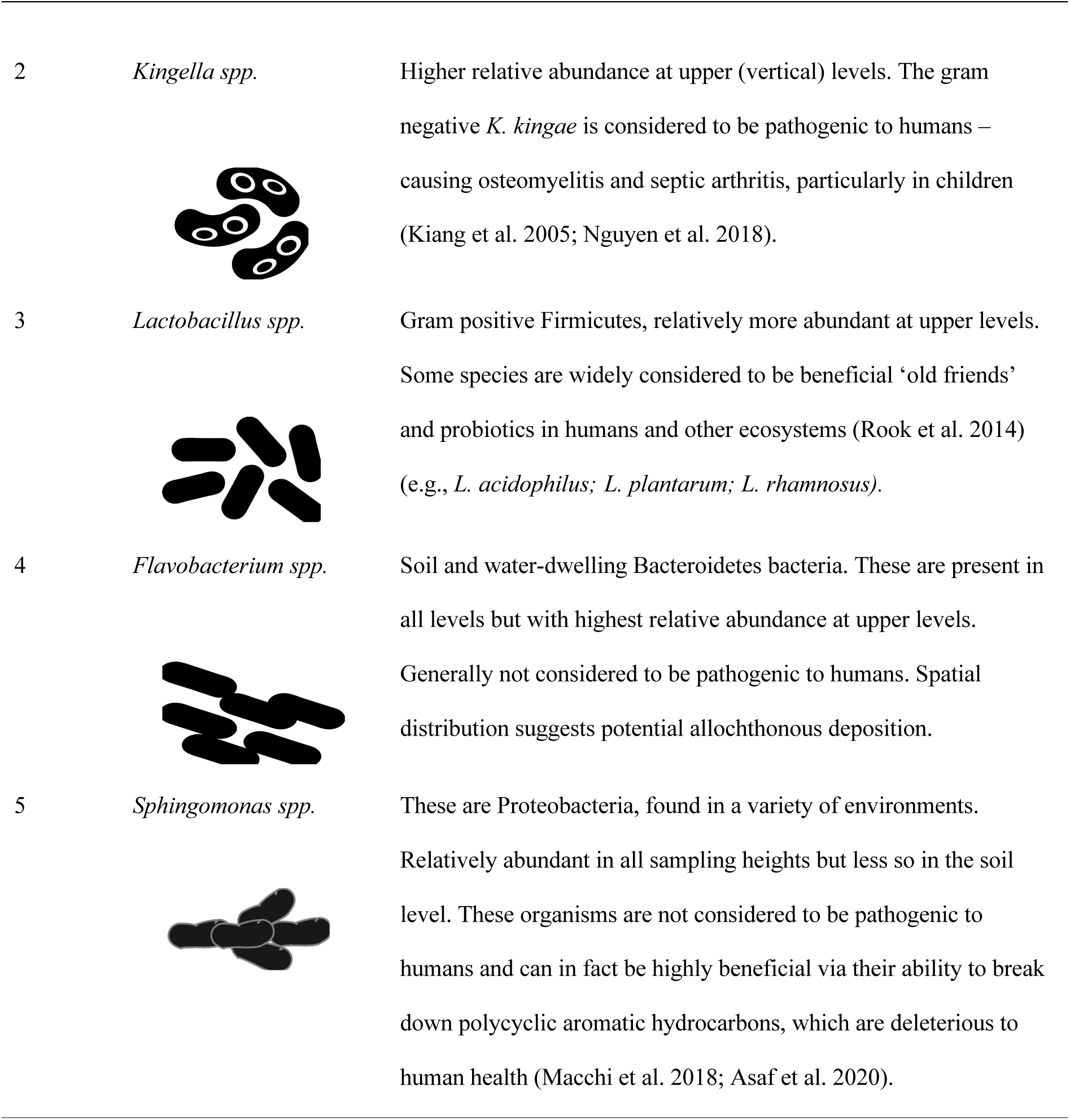
Notable taxa (OTUs at the genus level) identified during the examination for bacterial relative abundance – based on mean relative abundance (>1%) for each sampling height. These taxa may have important public health implications as highlighted in the third column.

| Number | Notable taxa | Potential public health implication |
| --- | --- | --- |
| 1 | *Streptomyces spp.* | These Actinobacteria are relatively more abundant at lower (vertically) sampling levels. They are soil-associated but also considered to be ‘old friends’ with potential beneficial implications for human health (Bolourian and Mojtahedi, 2018). |
| 2 | *Kingella spp.* | Higher relative abundance at upper (vertical) levels. The gram negative *K. kingae* is considered to be pathogenic to humans – causing osteomyelitis and septic arthritis, particularly in children (Kiang et al. 2005; Nguyen et al. 2018). |
| 3 | *Lactobacillus spp.* | Gram positive Firmicutes, relatively more abundant at upper levels. Some species are widely considered to be beneficial ‘old friends’ and probiotics in humans and other ecosystems (Rook et al. 2014) (e.g., *L. acidophilus*; *L. plantarum*; *L. rhamnosus*). |
| 4 | *Flavobacterium spp.* | Soil and water-dwelling Bacteroidetes bacteria. These are present in all levels but with highest relative abundance at upper levels. Generally not considered to be pathogenic to humans. Spatial distribution suggests potential allochthonous deposition. |
| 5 | *Sphingomonas spp.* | These are Proteobacteria, found in a variety of environments. Relatively abundant in all sampling heights but less so in the soil level. These organisms are not considered to be pathogenic to humans and can in fact be highly beneficial via their ability to break down polycyclic aromatic hydrocarbons, which are deleterious to human health (Macchi et al. 2018; Asaf et al. 2020). |

## 4 Discussion

### 4.1. Vertical stratification of aerobiome alpha diversity

Here we show that vertical stratification of aerobiome alpha diversity occurred. This transpired as a significant association in the reduction of bacterial alpha diversity as height increased (i.e., between the ground surface level and two vertical meters of the air column). When considering all sampling heights, alpha diversity reduced with greater height. This vertical stratification in alpha diversity was neither spatially (i.e., site specific) or temporally dependent. The strength of the negative relationship between alpha diversity and height increased when we merged lower sampling heights (0.0m with 0.5m) and the upper sampling heights (1.0m with 2.0m). This implies that the required spatial frequency to elucidate vertical stratification in alpha diversity––specifically, five sampling heights across a 2 m vertical transect––may have been overestimated. However, several omissions in the lower sampling heights due to failure to reach minimum DNA concentrations could have affected the strength of this association.

The decay in observed alpha diversity as height increased could be the result of increasing distance from the primary source, that is, potentially the soil. It is widely accepted that soil represents one of the most microbially-diverse terrestrial habitats (Briones, 2014; Bender et al. 2016; Dumbrell, 2019; Zhu et al. 2019). Therefore, it seems reasonable to suggest that lower sampling heights may possess a higher level of microbial diversity as they are closer to a potentially greater concentration of microbiota. We observed that a greater proportion of bacteria taxa found in the lower sampling heights (compared to the upper sampling heights) were also present in the soil samples, both at genus and family levels. Together, these results suggest that soil does appear to play a key role in supplementing the local aerobiome, particularly at lower heights.

The presence of vertical stratification of bacterial diversity in the aerobiome could have important implications for human health. Indeed, exposure to environmental microbes is thought to prime and ‘educate’ the immune system (Belkaid and Hand, 2014; Hanski, 2014; Minchim et al. 2020) particularly in early life, and a recent mouse study suggests that exposure to environmental microbes such as the butyrate-producer *Kineothrix alysoides* could also have anxiolytic (anxiety-reducing) effects (Liddicoat et al. 2019). The vertical stratification concept could also be important for exposome researchers, who investigate the types and methods of exposures to both endogenous and exogenous chemical composites (including microbes and their biological compounds across the life-course) (Escher et al. 2017; Daiber et al. 2019; McCall et al. 2019). The presence of vertical stratification implies that the potential for exposure to environmental microbial diversity may differ throughout the human life-course due to age and gender differences in height, activity types, and methods of motion.

### 4.2. Vertical stratification of aerobiome beta diversity

We also showed vertical stratification of aerobiome beta diversity, where sampling height explained 22% of the variation in environmental microbiota when all sampling heights were included. This was corroborated by the analysis of equality of taxonomic proportions between the air and the soil samples. As mentioned, the proportion of bacterial taxa from the air samples that were also present in the soil decreased as altitude increased. This provides preliminary evidence that soil has a stronger influence on aerobiome composition at lower heights and allochthonous sources make a key contribution to the aerobiome higher up.

It is likely that distance to source makes a key contribution to aerobiome vertical stratification. However, there may be other important biophysical driving factors. For example, the size range of bacterial cells can vary by eight orders of magnitude (from 0.013 μm to 750 μm) (Levin and Angert, 2015). However, many bacteria are thought to occur in the 0.3-5 μm range (Schaechter, 2016). Bacteria can also nucleate and exist as ‘clumps’ or adhere to larger suspended particles, thus altering their net particle size that would influence their fluid dynamics (Tham and Zuraimi, 2005; Haas et al. 2013; Gong et al. 2020). Airborne bacterial concentrations can be influenced by several factors including ambient temperature, humidity, wind dynamics and PM concentrations (Gong et al. 2020), and these factors could also play important roles in vertical stratification, and warrant further research.

Vertical stratification in bacterial *beta* diversity could also have important implications for public health. For example, our results point to intriguing questions such as: (a) are there significant and consistent differences in beneficial and pathogenic bacterial assemblages at different altitudes in the aerobiome? (b) does this significantly affect exposure and colonisation in humans across the life-course? (c) what are the downstream health implications of this, if any? We provide a preliminary contribution towards answering question (a), as discussed in the following section.

### 4.3. Relative abundances and notable taxa

Following the analyses of relative abundances, the dominant taxa in the soil and lower sampling heights were found to be Actinobacteria, and the dominant taxa in the upper sampling heights were Proteobacteria. This is not surprising given that a large proportion of terrestrial Actinobacteria are soil-dwelling organisms (Barka et al. 2016; Zhang et al. 2019), and both phyla are amongst the largest in the bacterial domain (Verma et al. 2013; Polkade et al. 2016; Rizzatti et al. 2017). Other studies have shown similar dominant roles for these phyla in the aerobiome (Arfken et al. 2015; Maki et al. 2017; Li et al. 2018), but vertical stratification has not, to our knowledge, been explored.

We identified a number of notable dominant taxa at the genus-level, including: *Streptomyces, Kingella, Lactobacillus, Flavobacterium*, and *Sphingomonas*. With the exception of *Flavobacterium*, species in these genera are known to have either beneficial or pathogenic impacts on human health. For example, the Actinobacteria *Streptomyces spp*., is considered to be a microbial ‘old friend’ and potentially beneficial to human health via production and regulation of anti-proliferative, anti-inflammatory and antibiotic compounds (Bolourian and Mojtahedi, 2018; Nguyen et al. 2020). This genus had higher relative abundance at lower sampling heights. On the other hand, members of the *Kingella* genus such as *K. kingae* are considered to be pathogenic to humans, for example––causing debilitating conditions such as osteomyelitis and septic arthritis, particularly in children (Kiang et al. 2005; Nguyen et al. 2018; Ingersol et al. 2019). These findings warrant further research––because if consistent across time and space, the spatial and compositional differences in microbiota have the potential to be important considerations for public health through the modulation of exposure.

## 5 Limitations

As a proof of concept study, we have demonstrated, for the first time, the presence of vertical stratification of microbial alpha and beta diversity at lower levels of the biosphere (ground level to 2.0 m high). However, we have not established the generalisability of our findings with a large number of replicates in different environments. Further, following the DNA extraction process, three samples (each at SC03 0.0 m) failed to reach sufficient DNA concentrations to enable PCR and sequencing, which may have affected the vertical stratification relationship––we can only speculate that the relationship would have been stronger with their inclusion. There are many sensitive variables involved with processing low biomass samples (Eisenhofer et al. 2019; McArdle and Kaforou, 2020) and perhaps even more stringent workflows are required for passive sampling.

## 6 Conclusions

We provide support for the presence of aerobiome vertical stratification in bacterial diversity (alpha and beta), and demonstrate that significant spatial differences in known pathogenic and beneficial bacterial taxa occur. Although the need to promote healthy ecosystems and understand environmental microbial exposures has always been important, in light of the COVID-19 pandemic, it is now justifiably at the forefront of many public health agendas worldwide. As discussed, there is growing evidence to suggest that exposure to the microbiome in biodiverse green spaces contributes towards ‘educating’ the immune system (Rook et al. 2003; Rook et al. 2013; Arleevskaya et al. 2019; Liddicoat et al. 2020). Furthermore, the microbiome is thought to support the immune system’s defensive role against pathogens, and prevent hyper-inflammatory responses and metabolic dysregulation–– risk factors for severe COVID-19 (Torres et al. 2019; Guo et al. 2020). Gaining a greater understanding of the transmission routes and physical factors (such as the vertical differential) affecting our exposure to environmental microbiomes––including both beneficial and pathogenic species––is likely to play an increasingly important role in the health sciences.

Strategies to explicitly consider the microbiome as part of health-promoting urban green spaces have recently been proposed, such as Microbiome-Inspired Green Infrastructure (MIGI) (Robinson et al. 2018; Watkins et al. 2020). Further exploration of aerobiome vertical stratification could make an important contribution to this approach. For example, there could be value in determining whether different habitats and vegetation management regimes impact vertical stratification in urban green spaces, and elucidating the downstream health effects on urban dwellers. Building on our findings––that vertical stratification did occur in an urban green space aerobiome––has the potential to inform future exposome research, urban biodiversity management, and disease prevention strategies.

## Conflicts of intrest

*The authors declare that the research was conducted in the absence of any commercial or financial relationships that could be construed as a potential conflict of interest*.

## Author Contributions

J.M.R and M.F.B contributed to the conception and design of the study; J.M.R and C.C.D conducted the field and lab work; J.M.R and C.L conducted the bioinformatics and data analysis; J.M.R wrote the manuscript; J.M.R produced the figures and data visualisations; J.M.R, C.C.D, C.L, P.W, R.C, and M.F.B contributed to manuscript internal critical review process and revisions. All authors read and approved the submitted version.

## Data Accessibility Statement

All data and code used in this study are available on the *UK Data Service ReShare* at https://reshare.ukdataservice.ac.uk; Data Collection #854411. All 16S rRNA gene sequences have been deposited in the European Nucleotide Archive (accession no. ERC000025).

## Acknowledgements

We acknowledge and pay our respects to the Kaurna people, the traditional custodians whose ancestral lands we conducted the research on for this study. We also acknowledge the City of Adelaide who helped to facilitate this study through time and resources.

